## Supplementary figure 1 for "VSTM1-v2 does not drive human Th17 cell differentiation"

### Supplementary Fig 1) VSTM1-v2 does not bind to leukocytes

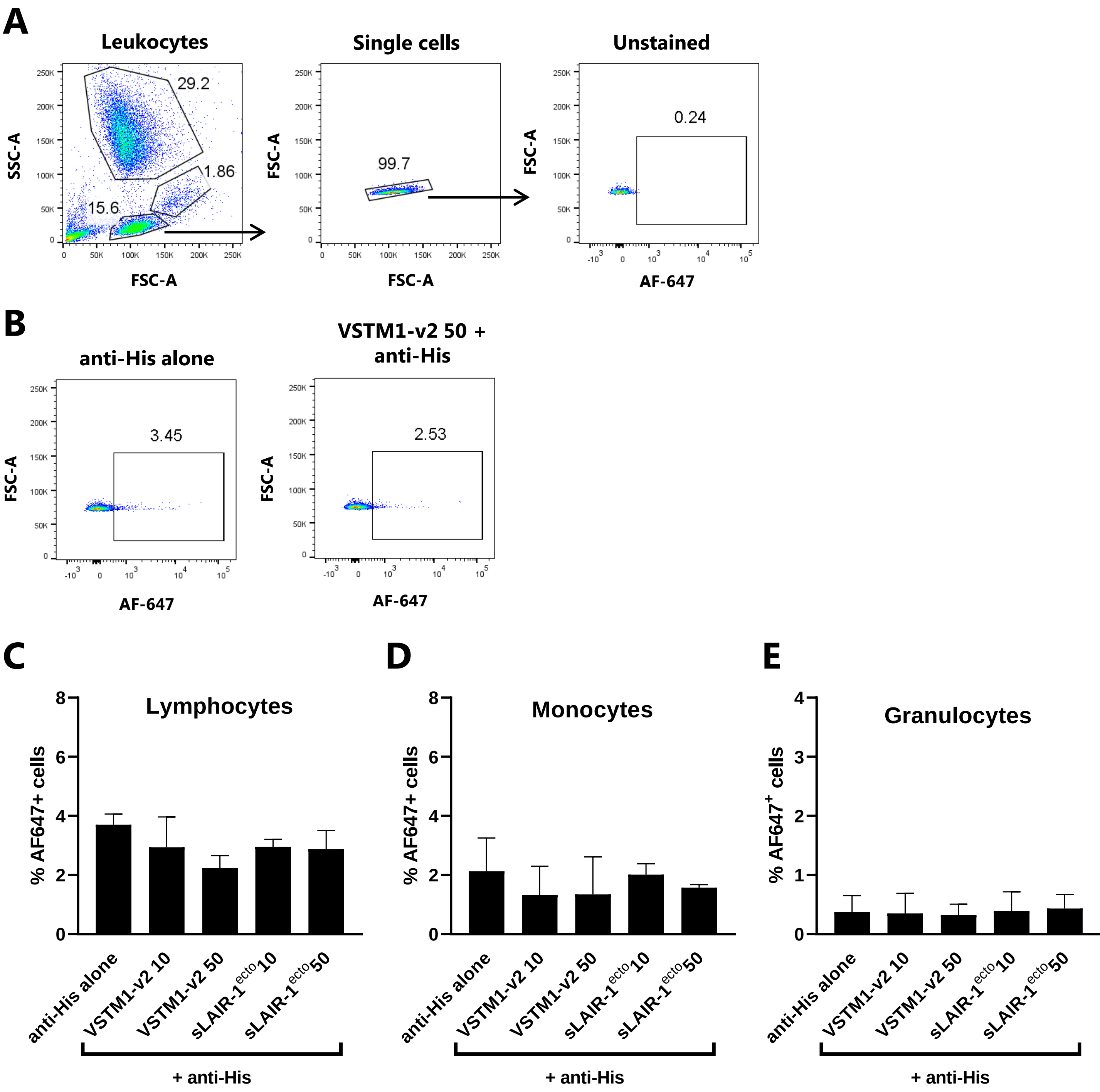


**Supplementary Fig 1. VSTM1-v2 does not bind to leukocytes.**
Erythrocyte-lysed whole blood was incubated with His-tagged VSTM1-v2 or sLAIR-1ecto (10-50 µg/mL), followed by detection with anti-His-AF647. (A) Lymphocytes (lower population), monocytes (middle population) and granulocytes (upper population) were gated based on forward scatter (FSC) and sideward scatter (SSC), followed by gating on single cells and AF-647+ cells. (B) Shown are representative dot plots of lymphocytes stained with 50 µg/mL VSTM1-v2 and/or anti-His-AF647. (C-E) The graphs show the quantification of the percentage of AF647+ cells of lymphocytes (C), monocytes (D), or granulocytes (E). Data are shown as mean ± SD of two donors.
